# A luciferase-based assay identifies niclosamide derivatives antagonizing Mcl-1 through post-translational down-regulation

**DOI:** 10.1101/665505

**Authors:** Qiang Liu, Jennifer M. Atkinson, Melat T. Gebru, Kristen Clements, George L. Moldovan, Hong-Gang Wang

## Abstract

The up-regulation of Mcl-1 expression is a major mechanism of cancer cell survival and therapy resistance. However, the underlying molecular mechanism remains incompletely understood, limiting the number of druggable approaches to selectively inhibit Mcl-1 function. In this study, we designed and employed a novel mechanistic high-throughput screening system to selectively uncover post-translational modulators of Mcl-1. We generated a cell-based high-throughput screening assay in which myeloid leukemia K562 cells constitutively express Mcl-1 or Bcl-xL fused with luciferase (Luc-Mcl-1 or Luc-Bcl-xL, respectively) under a viral promoter. 1,650 bioactive compounds were screened for their ability to selectively induce Mcl-1 down-regulation in a 2-hour assay. A family of niclosamide derivatives were eventually identified for their remarkable ability to decrease Mcl-1 protein stability, exemplified by N007. These salicylate derivatives did not alter Mcl-1 mRNA levels, but selectively induced proteasome-dependent Mcl-1 down-regulation independent of Noxa, Mule, or GSK3β. We also demonstrate that N007 potently induced cell death in leukemia cell lines, including those resistant to Bcl-2 inhibitors. Our work highlights the versatility of the mechanistic high-throughput screening approach as a valuable tool in identifying novel agents with the ability to down-regulate proteins crucial to human diseases.

## 1. Introduction

The evasion of apoptosis is a frequent feature of malignancies, and the dysregulation of B-cell lymphoma-2 (Bcl-2) family proteins through the up-regulation of Bcl-2, Bcl-xL or Mcl-1 is a major pathway by which this occurs [1,2]. The up-regulation of and functional dependence on anti-apoptotic Bcl-2 family proteins in cancer has led to the development of a number of small molecular antagonists of Bcl-2 family proteins. Such efforts have primarily focused on fragment-based approaches to design highly-specific protein-protein inhibitors that directly disrupt the interaction between anti-apoptotic and pro-apoptotic Bcl-2 family members through the BH3 domain [3]. At the forefront of such efforts is venetoclax (ABT-199), a small molecule that selectively binds to Bcl-2, but not Bcl-xL or Mcl-1 [4]. Venetoclax is highly efficacious in Bcl-2-dependent malignancies, but is ineffective in cancers dependent on Bcl-xL or Mcl-1.[5,6] Highly potent Bcl-xL inhibitors have also been developed, but clinical use is limited due to dose-limiting thrombocytopenia [7,8]. Given that numerous cancers have been shown to up-regulate and depend on Mcl-1 for survival and that Mcl-1 overexpression confers resistance to Bcl-2 inhibitors and other widely used anticancer therapies such as paclitaxel, vincristine and gemcitabine, selective Mcl-1 inhibitors are highly sought after [9,10].

A number of approaches have been undertaken to identify small molecule antagonists of Mcl-1. Traditionally, Mcl-1 antagonists have been developed by targeting up-stream kinases which regulate Mcl-1 transcription, of which sorafenib and dinaciclib are notable examples [11–15]. Fragment-based drug design has yielded BH3-domain mimetics that demonstrate high in vitro potency and selectivity towards Mcl-1. A potent and selective Mcl-1 inhibitor with in vivo activity, S63845, was recently disclosed and several other BH3 mimetic Mcl-1 inhibitors are under development. S63845 has been shown to stabilize Mcl-1 protein levels [16].

Alternative approaches to target Mcl-1, particularly small molecule post-translational modulation of Mcl-1, have remained relatively unexplored. The natural product maritoclax has now been described by multiple groups to induce Mcl-1-dependent cytotoxicity through post-translational modulation of Mcl-1 [15,17–20]. Unlike previously characterized inhibitors of Mcl-1, maritoclax does not alter Mcl-1 transcription, and does not function as a BH3 mimetic. Rather, maritoclax enhances the recruitment of Mcl-1 to the proteasome to induce its down-regulation [17,19]. Recently, a druggable motif on Mcl-1 has been described to regulate its ubiquitination and recruitment to the proteasome [21]. Together, this data suggests that the post-translational antagonism of Mcl-1 is a feasible but relatively unexplored approach to elucidate novel small molecule inhibitors of Mcl-1 function.

To this end, we designed and validated a robust high-throughput screening platform to identify bioactive compounds that antagonize Mcl-1 through post-translational modulation. Using our novel screening platform, we have identified four compounds which induce Mcl-1 proteasome dependent down regulation: geldanamycin, niclosamide, PP121, and WP1130. As proof-of-principle, we performed structure-analysis studies on niclosamide and uncovered potent salicylamide derivatives which post-translationally destabilize Mcl-1.

## 2. Materials and Methods

### Plasmids, antibodies, and compounds

The MSCV-Luc-Mcl-1 and MSCV-Luc-Bcl-xL vectors were generated by subcloning luciferase and puromycin from the MSCV-luciferase-IRES-YFP vector (Dr. Gerard Grosveld, St. Jude Children’s Research Hospital) into the MSCV-Mcl-1-IRES-Bim or MSCV-Bcl-xL-IRES-Bim vector [17] and deleting the IRES-Bim.

Antibodies were obtained from the following sources: Mcl-1 [22], Bcl-2 [22], Bcl-xL (Cell Signaling #2764), Bim (Sigma B7929), Bax (Santa Cruz sc-493), Mule/HUWE1 (Bethyl A33-486A), GSK3 (Cell Signaling #5676), Noxa (Thermo Fisher MA1-41000), and Actin (Sigma A2228). Immunoblotting was performed as previously described following cell lysis in RIPA buffer [17] with the exception of experiments involving Mule^KO^ HeLa cells, where lysates were prepared as outlined previously [23]. Blots were scanned on the Licor Odyssey CLx and quantified using ImageStudio version 5.0.

The following compounds were used in this study: the bioactive compound library was ordered from Selleck Chemicals (L1700); maritoclax and KS18 were synthesized as previously described [17,20], cycloheximide (Sigma C7698); geldanamycin (LKT Labs G1646); niclosamide (Sigma N3510); PP121 (Selleck S2622); WP1130 (Selleck S2243); MG132 (LKT Labs M2400); Z-VAD-FMK (Enzo ALX-260-020-M001); niclosamide analogues N001-N021 were ordered from Mcule.

### Cell lines and transfection

K562, Raji, KG-1, U937, H929, and HL60, cells were obtained from ATCC (CCL-243, CCL-86, CCL-246, CRL-1593, CRL-9068, and CCL-240 respectively) and cultured in IMDM medium (Corning 10-016-CV) with 10% FBS (Sigma F2442) and 1% antibiotic-antimycotic solution (Corning 30-004-CI). NSCLC cell lines H23 and H460 were maintained in RPMI-1640 medium (Corning 10-040-CI) with 10% FBS and 1% antibiotic-antimycotic solution. Mule^KO^ HeLa cells were generated as previously described [23]. Noxa^KO^ HCT116 cells and the matching parental controls were kindly received from Dr. Jian Yu (University of Pittsburgh). GSK3B^KO^ MEF cells and matching parental controls were kindly provided by Dr. James R. Woodgett (Lunenfeld-Tanenbaum Research Institute) [24].

Luc-Mcl-1/K562 and Luc-Bcl-xL/K562 cells were generated using the previously described retrovirus transduction protocol [17]. Stable cell lines were generated through puromycin selection and maintained in IMDM medium with 10% FBS and 1% antibiotic-antimycotic solution.

### Luminescence activity assays in Luc-Mcl-1/K562 and Luc-Bcl-xL/K562 cells

Luc-Mcl-1/K562 and Luc-Bcl-xL/K562 cells were seeded at 1 × 10^6^ cells/mL in 96- or 384-well plates in IMDM medium with 2% FBS and 1% antibiotic-antimycotic solution with compounds in less than 0.5% DMSO unless otherwise stated. Luciferase reagent (Promega One-Glo, E6120) was added to cells after compound treatment and incubated for 10 minutes according to manufacturer recommendations. Luminescence was measured with one hour of reagent addition on the BMG ClarioStar.

### High-throughput screen of bioactive compounds in Luc-Mcl-1 K562 cells

The Selleck Chemicals bioactive library was reformatted into a 384 well format using a Tecan Freedom Evo liquid handler. Luc-Mcl-1 cells were seeded into white 384 well plates at a density of 20,000 cells per well using a reagent dispenser (Biotek MultiFlo FX) and compounds added to cells using the Tecan Freedom Evo for a final concentration of 10µM in a 30µl volume, 1% DMSO. WP1130 was used as a positive control compound and to monitor for consistency across screening plates. Cells were incubated at 37°c, 5% CO_2_ for 90 minutes before being equilibrated to room temperature for 30 minutes and the luminescence assay performed as described above.

### Secondary screening of active compounds

Compounds which demonstrated a response greater than 2.5 standard deviations from the mean (65%) where selected as ‘hits’ from the primary screen and tested in dose response analysis in Luc-Mcl-1 and Luc-Bcl-xL/K562 cells. In secondary screening compounds were cherry picked from the drug screening plates and used for a 11-point dose response analysis. Dilutions (1:2) were performed using EP Motion (Eppendorf) and transferred to cells using the Tecan Freedom Evo. Final compound concentrations ranged from 10µM to 4.9nM in 1% DMSO with 20,000 cells per well. Compounds active in secondary screening were re-sourced and validated.

### Cell viability assays

Cells were plated at a density of 7,500 cells/30µl/well in white 384 well plates. After 24 hours of incubation, cells were treated with drug at the desired concentrations. 48 hours after compound addition, cell viability was measured using CellTiter-Glo (Promega G7571) according to manufacturer’s instructions.

### Real-time quantitative reverse-transcriptase PCR (qRT-PCR)

Sample preparation for qRT-PCR was performed as previously described [19]. qRT-PCR and data analysis were performed on the QuantStudio 12K Flex Real-Time PCR System (ThermoFisher Scientific). qRT-PCR was normalized using the geometric mean of the following internal control genes: ACTB, HMBS, SDHA, TBP, and B2M [25]. The following primers were used: sense 5’-GGACATCAAAAACGAAGACG-3’ and antisense 5’-GCAGCTTTCTTGGTTTATGG-3’ for MCL1; sense 5’-AGCTGGAAGTCGAGTGTGCT-3’ and antisense 5’-TCCTGAGCAGAAGAGTTTGGA-3’ for PMAIP1; sense 5’-CGGGAAATCGTGCGTGACATTAAG-3’ and antisense 5’-TGATCTCCTTCTGCATCCTGTCGG-3’ for ACTB; sense 5’-ACCAAGGAGCTTGAACATGC-3’ and antisense 5’-GAAAGACAACAGCATCATGAG-3’ for HMBS; sense 5’-TGGGAACAAGAGGGCATCTG-3’ and antisense 5’-CCACCACTGCATCAAATTCATG-3’ for SDHA; sense 5’-TGCACAGGAGCCAAGAGTGAA-3’ and antisense 5’-CACATCACAGCTCCCCACCA-3’ for TBP; sense 5’-CTCCGTGGCCTTAGCTGTG-3’ and antisense 5’-TTTGGAGTACGCTGGATAGCCT-3’ for B2M.

### Statistics

All statistical analyses were performed using GraphPad Prism version 6.00 for Windows (GraphPad Software, www.graphpad.com). EC_50_ calculations for viability were calculated through nonlinear regression with normalized data assuming variable slope. Other statistical tests were performed as indicated.

## 3. Results

### 3.1. Luciferase-tagged Mcl-1 and Bcl-xL mimic endogenous protein turnover dynamics in K562 cells

In order to specifically assay the post-translational regulation of Mcl-1, we constructed a fusion protein containing firefly luciferase fused to either Mcl-1 or Bcl-xL (Luc-Mcl-1 and Luc Bcl-xL, respectively) under the control of the murine stem cell virus (MSCV) promoter (**Figure 1A**). In order to confirm that the luciferase tag did not affect the fusion protein’s stability, we compared the protein half-lives of tagged Mcl-1 and Bcl-xL to their endogenous counterparts in K562 cells. We confirmed that the protein half-lives between the tagged and endogenous protein pairs did not differ in the presence of cycloheximide, an inhibitor of protein translation (**Figure 1B**). Similarly, the luminescent activity of K562 cells expressing Luc-Mcl-1 (Luc-Mcl-1/K562) or Luc-Bcl-xL (Luc-Bcl-xL/K562) closely correlated with the respective protein levels and published protein half-life values (**Figure 1C**).

**Figure 1.**
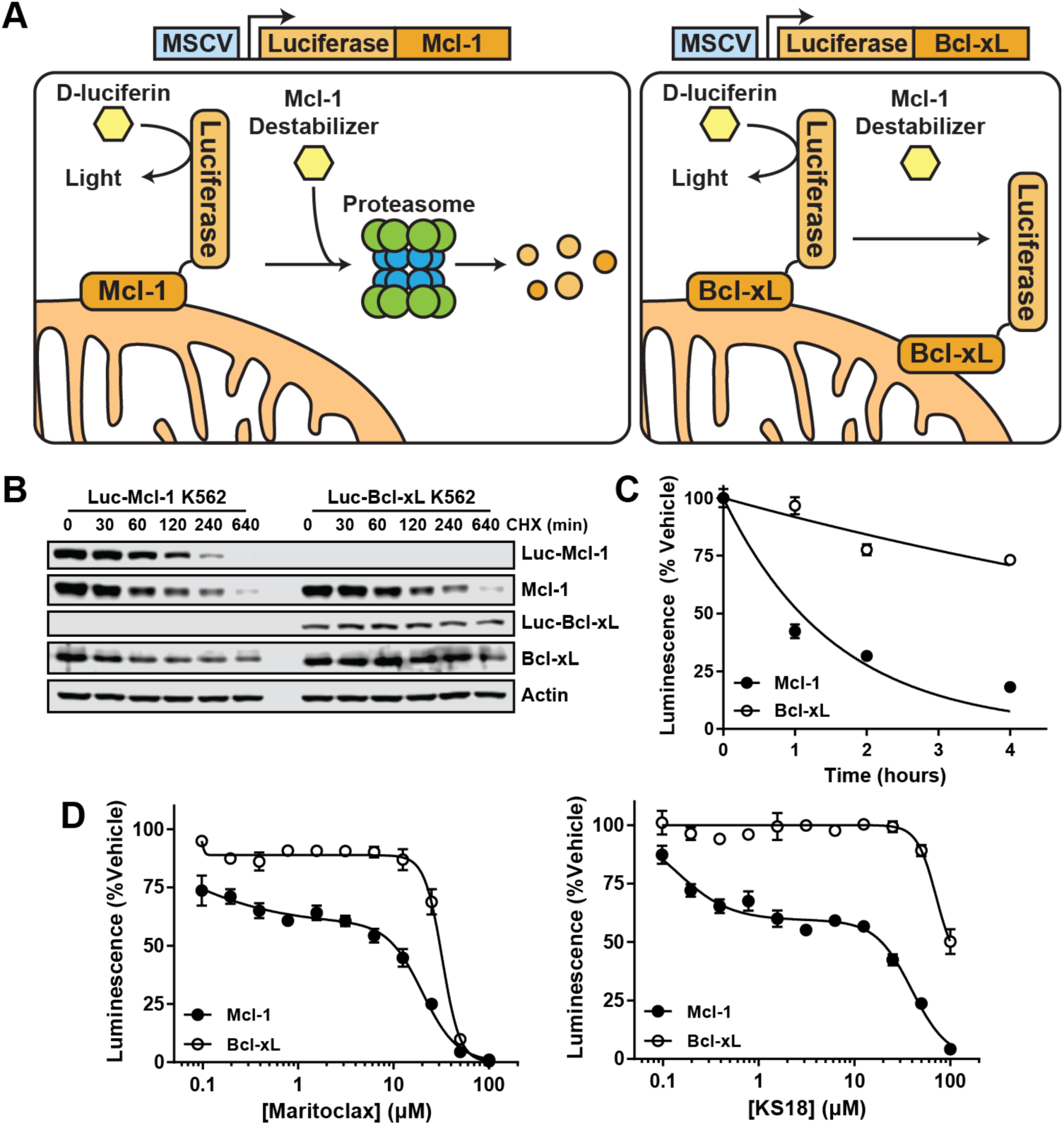
Luciferase-tagged Mcl-1 and Bcl-xL mimic endogenous protein dynamics. (**a**) Schematic demonstrating the principle by which luciferase-tagged Mcl-1 might be used to uncover post-translational modulators of Mcl-1, with luciferase-tagged Bcl-xL as control. (**b**) K562 cells stably expressing luciferase-tagged Mcl-1 (Luc-Mcl-1) or Luc-Bcl-xL (Luc-Bcl-xL) were treated with 10µg/mL cycloheximide (CHX) for the indicated times and subjected to immunoblotting. Representative data of two independent experiments is shown. (**c**) Luc-Mcl-1/K562 and Luc-Bcl-xL/K562 cells were treated with 10µg/mL cycloheximide, and luminescence was measured at the indicated times after cycloheximide addition and normalized to time 0 (T_0_). Error bars represent the standard deviation of three replicates. (**d**) Luc-Mcl-1/K562 and Luc-Bcl-xL/K562 were treated with the indicated concentrations of maritoclax and KS18, and luminescence was measured two hours. Representative data of two independent experiments is shown. Error bars represent that standard deviation of four replicates.

Maritoclax and KS18 have previously been demonstrated to induce Mcl-1 degradation by the proteasome [19,20]. Thus, to confirm whether Luc-Mcl-1 is sensitive to post-translational modulation in response to small molecules, Luc-Mcl-1/K562 and Luc-Bcl-xL/K562 were treated with maritoclax and KS18. Both compounds were observed to inhibit Luc-Mcl-1 over Luc-Bcl-xL activity with potencies that closely match previously published IC_50_ values (**Figure 1D**) [19,20]. Together, this data indicates that the Luc-Mcl-1/K562 and Luc-Bcl-xL/K562 cell lines are simple and robust systems in which potential post-translational modulators can be assayed on a high throughput scale.

### 3.2. A high-throughput screen identifies bioactive compounds that selectively modulate Mcl-1

A screening workflow was established to identify and validate compounds which may selectively destabilize Mcl-1 through post-translational modulation (**Figure 2A**). In the primary screen, 1650 bioactive compounds were screened in Luc-Mcl-1/K562 cells. Here, 64 compounds that displayed activity greater than 2.5-fold of the standard deviation from the mean were chosen for the secondary dose-response screen in both Luc-Mcl-1/K562 and Luc-Bcl-xL/K562 cells (**Figure 2B**). Following secondary and tertiary screens, eight compounds were determined to demonstrate some degree of selective inhibition towards Luc-Mcl-1 over Luc-Bcl-xL reporter (**Figure 2A**). Four compounds were eventually validated for their capacity to down-regulate endogenous Mcl-1 protein in Raji cells: geldanamycin, niclosamide, PP121, and WP1130 (**Figure 2C,D**). Although WP1130 only displayed marginal selectivity towards Luc-Mcl-1, the compound was shown to be selective towards endogenous Mcl-1 over Bcl-2 and Bcl-xL, suggesting that WP1130 may destabilize luciferin-tagged proteins (**Figure 2C,D**). Additionally, these four compounds were tested in Raji cells for their capacity to suppress the transcription of Mcl-1 mRNA. Compared to dinaciclib, a cyclin-dependent kinase (CDK) inhibitor that potently suppresses Mcl-1 transcription, none of the four lead compounds suppressed Mcl-1 transcription (**Figure 2E**). Thus, four compounds were validated to down-regulate Mcl-1 without suppressing its mRNA transcription: geldanamycin, niclosamide, PP121, and WP1130.

**Figure 2.**
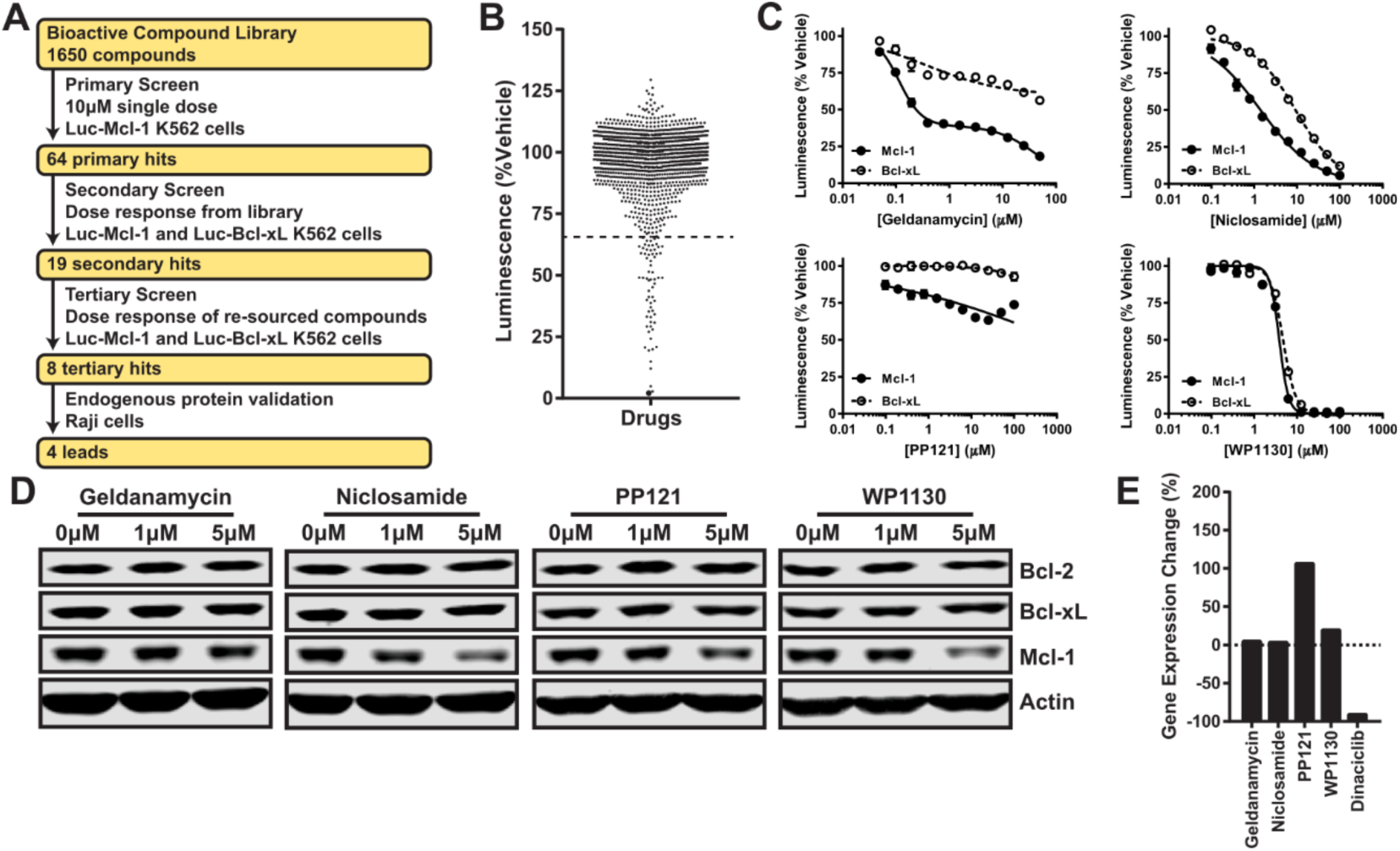
High-throughput screen of bioactive compounds reveal small molecules that down-regulate Mcl-1 protein levels without suppressing transcription. (**a**) Schematic showing the workflow followed by the high-throughput screen. (**b**) Luc-Mcl-1/K562 cells were screened with 1650 bioactive compounds over 2 hours, and the responses are shown as percent inhibition of Luc-Mcl-1 luminescence. Dotted line marks the response that is 2.5-fold standard deviations away from the median. (**c**) Luc-Mcl-1/K562 and Luc-Bcl-xL/K562 cells were treated with the indicated concentrations of the compounds for 2 hours, followed by luminescence measurement. Error bars represent the standard deviation of three independent experiments. (**d**) Raji cells were treated with the indicated concentrations of compounds over 8 hours and subjected to immunoblotting. (**e**) Raji cells were treated with 5µM of each compound, and 100nM of dinaciclib, over 4 hours and subjected to qRT-PCR analysis.

### 3.3. Salicylamide derivatives selectively down-regulate Mcl-1

Geldanamycin, PP121, and WP1130 have known targets and have been amply described in the literature for their antitumor potential [26–29]. Niclosamide is an anthelmintic that is only recently under investigation for a myriad of antitumor activities. Niclosamide has been described to antagonize STAT3 or Bcl-2 family proteins but its molecular mechanism has not been described [30–33]. Since niclosamide does not have a well-defined target, we chose to work up this compound as a novel antagonist of Mcl-1. In order to determine whether the chemical structure of niclosamide could be optimized to yield more effective small molecules with enhanced selectivity in antagonizing Mcl-1, we selected a series of niclosamide analogues with a salicylamide backbone and determined their potency in our Luc-Mcl-1 and Luc-Bcl-xL reporter assay (**Table 1**). We discovered a number of salicylamide derivatives that demonstrate significantly enhanced selectivity towards Luc-Mcl-1 compared to niclosamide itself (**Figure 3A**). Among the salicylamides tested, N007 exerted marked down-regulation for endogenous Mcl-1 protein (**Figure 3B**).

**Table 1.** Structures and activities of salicylamide derivatives tested in this study. IC_50_ values represent the concentration for which half of the luciferase activity was inhibited in Luc-Mcl-1/K562 or Luc-Bcl-xL/K562 cells. IC_50_ values are the average of two independent repeats.

| Compound | R1 | R2 | R3 | R4 | R5 | R6 | R7 | R8 | R9 | Luc-Mcl-1 | Luc-Bcl-xL |
| --- | --- | --- | --- | --- | --- | --- | --- | --- | --- | --- | --- |
| Niclosamide | - Cl | - H | - H | - H | - H | - H | -NO<sub>2</sub> | - H | - Cl | 1.37 | 7.51 |
| N001 | - Cl | - H | - H | - H | - H | - H | - H | - Cl | - H | 9.37 | > 100 |
| N002 | - H | - H | - H | - H | - H | - H | - H | - H | - Cl | > 100 | > 100 |
| N003 | - Cl | - H | - H | - H | - H | - H | - NHCOCH(CH<sub>3</sub>)<sub>2</sub> | - H | - H | > 100 | > 100 |
| N004 | - H | - H | - H | - H | - H | - H | - NHCOCH<sub>3</sub> | - Cl | - H | > 100 | > 100 |
| N005 | - Cl | - H | - H | - H | - H | - H | - H | - Cl | - Cl | 10.47 | > 100 |
| N006 | - Cl | - H | - H | - H | - H | - H | - H | - NHCOCH<sub>3</sub> | - H | > 100 | > 100 |
| N007 | - Cl | - H | - H | - H | - H | - H | - Cl | - Cl | - H | 4.77 | > 100 |
| N008 | - Cl | - H | - H | - H | - H | - H | - H | - H | - H | 37.84 | > 100 |
| N009 | - Cl | - H | - H | - H | - H | - H | - Cl | - H | - H | 6.55 | > 100 |
| N010 | - Cl | - H | - H | - H | - H | - H | - NH<sub>2</sub> | - H | - Cl | > 100 | > 100 |
| N011 | - Br | - H | - H | - H | - H | - H | - NO<sub>2</sub> | - H | - Cl | 0.66 | 5.87 |
| N012 | - H | - H | - H | - H | - H | - Cl | - H | - H | - Cl | > 100 | > 100 |
| N013 | - Cl | - H | - H | -CH<sub>3</sub> | - H | - H | - H | - Cl | - Cl | > 100 | > 100 |
| N014 | - Cl | - H | - H | - H | - H | - H | - CH<sub>3</sub> | - H | - H | 16.56 | > 100 |
| N015 | - Cl | - H | - NO<sub>2</sub> | - H | - H | - Cl | - H | - H | - Cl | 19.54 | > 100 |
| N016 | - Cl | - C<sub>6</sub>H<sub>5</sub> | - C<sub>6</sub>H<sub>5</sub> | - H | - H | - H | - NO<sub>2</sub> | - H | - Cl | 5.31 | 51.17 |
| N017 | - Cl | - C<sub>6</sub>H<sub>5</sub> | - C<sub>6</sub>H<sub>5</sub> | - H | - H | - H | - H | - H | - Cl | > 100 | > 100 |
| N018 | - Cl | - H | - Cl | - H | - H | - H | - NO<sub>2</sub> | - H | - Cl | 9.06 | 59.29 |
| N019 | - Br | - H | - Br | - H | - H | - H | - NO<sub>2</sub> | - H | - Cl | 5.04 | 30.69 |
| N020 | - Cl | - H | - Cl | - H | - H | - H | - NHCOCH<sub>3</sub> | - H | - H | > 100 | > 100 |
| N021 | - Cl | - H | - H | - H | - Cl | - H | - NO<sub>2</sub> | - H | - Cl | 18.71 | > 100 |

**Figure 3.**
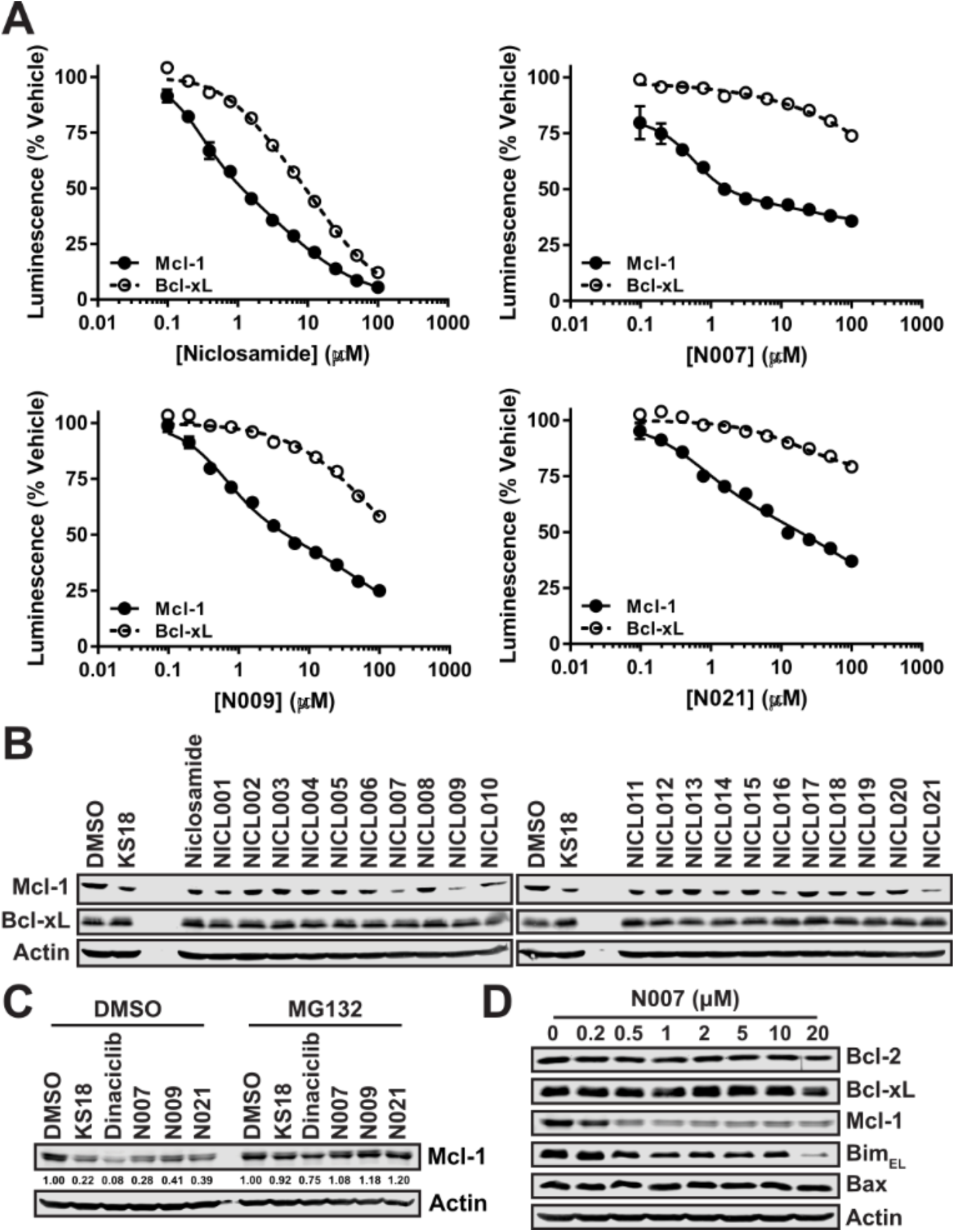
Niclosamide analogues selectively down-regulate Mcl-1. (**a**) Luc-Mcl-1/K562 and Luc-Bcl-xL/K562 cells were treated with the indicated concentrations of compounds over two hours, followed by luminescence measurement. Representative data of two independent experiments is shown. Error bars represent the standard deviation of four replicates. (**b**) Raji cells were treated with 5µM of the indicated compounds for 8 hours and subjected to immunoblotting. (**c**) Raji cells were treated with the following concentrations of compounds as indicated for 8 hours and subjected to immunoblotting: KS18, N007, N009, and N021 (5µM); dinaciclib (100nM); MG132 (10µM). Numbers indicate the quantified Mcl-1 protein levels relative to DMSO-treated control. (**d**) Raji cells were treated with the indicated concentrations of N007 over 24 hours and subjected to immunoblotting.

Interestingly, the chemical structure of N007 resembled the structure of pyoluteorins previously characterized to promote Mcl-1 proteasomal degradation, such as KS18 [20]. Thus we sought to determine whether these salicylamides could similarly induce the proteasomal degradation of Mcl-1. Expectedly, the treatment of MG132, a proteasome inhibitor, blocked the Mcl-1 degradation induced by KS18 but not dinaciclib (**Figure 3C**). MG132 was similarly able to completely block Mcl-1 down-regulation induced by active salicylamide derivatives including N007, N009, and N021. We confirmed that N007 indeed selectively down-regulated Mcl-1 protein levels among Bcl-2 family member proteins (**Figure 3D**). An apparent down-regulation of Bim_EL_ following N007 treatment might indicate the tandem degradation of the Mcl-1/Bim_EL_ complex. Given its potency and selectivity among the tested salicylamides, we selected N007 as the prototypical salicylamide for future studies as a novel antagonist of Mcl-1.

### 3.4. Proteasome-dependent degradation of Mcl-1 mediated by salicylamide derivatives is not dependent on Noxa, Mule, or GSK3B

A decrease in the post-translational protein stability manifests in a decrease in protein half-life. To ascertain if salicylamide derivatives such as N007 post-translationally destabilize Mcl-1, N007 was co-treated with cycloheximide over 4 hours. We determined that N007 significantly decreased the protein half-life of Mcl-1 (**Figure 4A,B**). Additionally, N007 was not found to decrease Mcl-1 transcription levels through qRT-PCR (**Figure 4C**). On the other hand, we observed a time-dependent increase in Mcl-1 transcription, which we speculate to be due to feedback mechanisms from Mcl-1 protein down-regulation.

**Figure 4.**
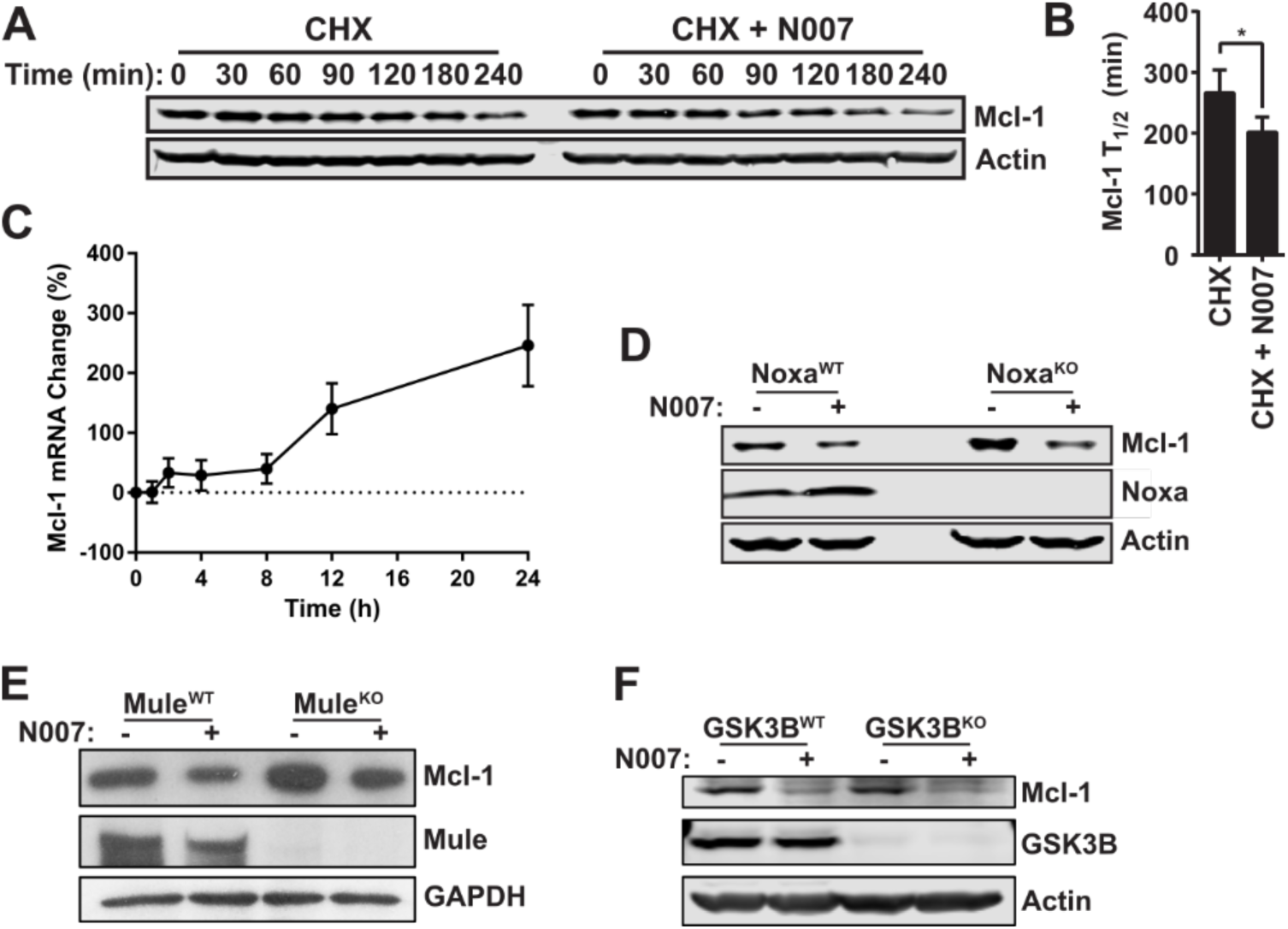
Niclosamide analogue post-translationally down-regulates Mcl-1 independent of Noxa, Mule, or GSK3B. (**a,b**) Raji cells were treated with 10µg/mL cycloheximide (CHX) with vehicle control or 5µM N007, and subjected to immunoblotting at the indicated times. The representative blots of three independent experiments are shown in (**a**). The quantified Mcl-1/Actin protein ratios are shown in (**b**). Error bars represent the standard deviation of three independent repeats. Statistics were calculated by Student’s T-test. \**p* < 0.05. (**c**) Raji cells were treated with 5µM of N007 and samples were collected for qRT-PCR analysis at the indicated times. The combined data of two independent experiments are shown, and error bars represent the standard deviation. (**d**) Parental HCT116 cells (Noxa^WT^) and HCT116 cells bearing homozygous Noxa deletions (Noxa^KO^) were treated with 5µM of N007 for 24 hours and subjected to immunoblotting. The representative blot of three independent repeats is shown. (**e**) Parental HeLa cells (Mule^WT^) and HeLa cells bearing homozygously deleted Mule (Mule^KO^) were treated with 5µM of N008 for 24 hours and subjected to immunoblotting. The representative of two independent experiments are shown. (**f**) Parental MEFs (GSK3B^WT^) or MEFs bearing homozygously deleted GSK3B (GSK3B^KO^) were treated with 5µM N007 for 24 hours and subjected to immunoblotting. The representative blot of three independent experiments is shown.

A number of proteins have been described to regulate the post-translational stability of Mcl-1, and we sought to determine if the observed proteasome dependent down-regulation of Mcl-1 was due to the effects of N007 on some of these key proteins. Among BH3-only Bcl-2 family proteins, Noxa is unique in its capacity to post-translationally destabilize Mcl-1 upon binding [34–36]. To this end, we used HCT116 cells bearing homozygously deleted Noxa (Noxa^KO^). Surprisingly, N007 significantly suppressed Mcl-1 protein levels despite the lack of Noxa, suggesting that Noxa does not have a functional role in Mcl-1 down-regulation induced by N007 (**Figure 4D**).

The proteasomal degradation of polyubiquitinated Mcl-1 contributes to its rapid protein turnover. A number of E3 ubiquitin ligases have thus been described to regulate Mcl-1 protein stability. Among these, the E3 ubiquitin ligase HUWE1, also known as Mcl-1 ubiquitin ligase E3 (Mule), has been described to play a major role in Mcl-1 ubiquitination [37]. Two other E3 ubiquitin ligases have also been described for Mcl-1: FBW7 and β-TrCP. Both of these E3 ubiquitin ligases are reported to be canonically dependent on GSK3B-mediated phosphorylation of Mcl-1 [38,39]. To determine the functional contribution of these E3 ubiquitin ligases to N007-mediated Mcl-1 suppression, we used cell lines bearing deletions in Mule (Mule^KO^) and GSK3B (GSK3B^KO^) [23,24]. We observe that N007 suppressed Mcl-1 in both of these knock-out cell lines, suggesting that down-regulation of Mcl-1 induced by salicylamide derivatives do not depend on these E3-ubiquitin ligases (**Figure 4E,F**).

### 3.5. Cancer cell lines undergo cell death following N007 treatment

In order to determine whether N007 was able to reduce cell viability, we treated a number of cancer cell lines with N007, as well as the Bcl-2 inhibitor ABT-199 and Bcl-2/Bcl-xL inhibitor ABT-737 [4,5]. The selected cancer cell lines have previously been characterized for their Bcl-2 family expression: The cell lines Kasumi-1, HL60, U937, H929, and H23 cells have been reported to be high in Mcl-1 expression [19,40,41], and KG-1 and H460 cells have been reported to be low in Mcl-1 and high in Bcl-2 or Bcl-xL [19,40]. We observed that N007 was active in reducing the viability of the tested cell lines at IC_50_ values closely matching to its IC_50_ for down-regulating Mcl-1 protein in our screening assay (**Table 2**). We also observed that N007 was active against cell lines that were resistant to ABT-199 and ABT-737. This data suggests that salicylamides can induce apoptosis in cancer cell lines and that Mcl-1 down-regulation may contribute to the observed cytotoxicity.

**Table 2.** The potency of the indicated compounds against a panel of cancer cell lines over 48 hours, expressed as the IC_50_ (µM).

|  | Kasumi-1 | KG-1 | U937 | HL60 | H929 | H23 | H460 |
| --- | --- | --- | --- | --- | --- | --- | --- |
| **N007** | 3.7 | 5.6 | 1.1 | 1.7 | 1.3 | 2.4 | 1.9 |
| **ABT-199** | 0.1 | 6.7 | 9.0 | 0.2 | > 10 | > 10 | > 10 |
| **ABT-737** | 0.1 | 1.5 | 3.9 | 0.2 | > 10 | > 10 | > 10 |

## 4. Discussion

The development of post-translational Mcl-1 inhibitors is an underexplored avenue in anti-cancer drug development. Here we describe the development of a high throughput screening system designed to identify small molecules which promote the selective degradation of Mcl-1 relative to Bcl-xL. Our studies identified salicylamide derivatives which act to promote proteasome-dependent Mcl-1 degradation independent of Noxa, Mule and GSK3β and promote cell death in cell lines that express elevated levels of Mcl-1 (**Fig. 4 and Table 2**).

Several small molecules specifically inhibit the apoptotic roles of Mcl-1 by targeting its BH3 protein binding domain [16,42–44]. These compounds protect Mcl-1 from proteolysis and in turn result in significant up-regulation of Mcl-1 protein levels. As Mcl-1 has been shown to play roles in cell cycle, chemotherapy-induced senescence, mitochondrial respiration, and calcium signaling independent of its BH3 domain [45–47], the precise effects of Mcl-1 protein up-regulating coincident with BH3 antagonism is unclear. It is likewise unknown whether Mcl-1 protein down-regulation would be more efficacious for cancer therapy compared to BH3 domain inhibition which results in stabilization of the protein.

To address this, we designed a mechanistic high-throughput screening approach that would elucidate Mcl-1 antagonists which decrease its protein expression. By placing the Mcl-1-luciferase fusion protein under a constitutive promoter, a compound library screened revealed a number of compounds that could selectively target Mcl-1 for post-translational down-regulation. This screening approach allowed us to elucidate four bioactive compounds which down-regulate Mcl-1 (**Figure 2**). WP1130 is an inhibitor of USP9X, a deubiquitinase for Mcl-1. Accordingly, WP1130 has been described to exert cytotoxicity by post-translationally down-regulating Mcl-1 [28]. Identifying WP1130 through our novel screening system serves as further evidence for the success of our screening design. PP121 is a multi-kinase inhibitor, but the compound has not been described in the context of Mcl-1. Nonetheless, several kinases have been described to modulate the post-translational stability of Mcl-1, and it is likely PP121 acts through one or more of these pathways to selectively degrade Mcl-1 [48]. Geldanamycin is an Hsp90 inhibitor, and Hsp90 inhibitors have recently been reported to modulate Mcl-1 thr006Fugh GSK3β and FBW7 [49]. Finally, we identified niclosamide, an anthelmintic agent, as a potent inhibitor of Mcl-1. Although niclosamide has been reported to be biologically active against various cancer cells, its primary mechanism of action is unknown [30–32,50].

SAR studies on niclosamide revealed a salicylamide pharmacophore that coincidentally resembles a series of Mcl-1 inhibitors previously characterized in our laboratory, such as maritoclax and KS18 [17,19,20]. Similar to maritoclax, active niclosamide analogues such as N007 selectively destabilized Mcl-1 post-translationally. Overall, we determined that N007 induces post-translational Mcl-1 destabilization independent of Mule and Noxa. Present data suggests that salicylamide derivatives behave through non-canonical mechanisms of Mcl-1 protein regulation, or through the recently reported E3 ligase for Mcl-1, MARCH5 [51]. Given the structural and mechanistic similarity between N007 and KS18, it is possible that these two compounds bind to a similar site on Mcl-1.

Here, we have validated a novel and robust screening approach to uncover post-translational modulators of Mcl-1, and characterized niclosamide as a novel Mcl-1 down-regulator. The fusion of reporter constructs to proteins of interest should facilitate the discovery of traditionally undruggable targets. This approach can also be utilized to screen novel regulators of protein stability, for instance by employing genome-wide siRNA or CRISPR/Cas9 screens. Our study collectively demonstrates the value of our novel screening platform, and identifies salicylamide derivatives as post-translational modifiers of Mcl-1 worthy of further study and validation.

## Funding

This work was supported by the Lois High Berstler Research Endowment Fund and the Four Diamonds Fund of the Pennsylvania State University College of Medicine.

## Acknowledgments

We thank Dr. Jian Yu from the University of Pittsburgh for generously providing the Noxa^KO^ HCT116 cells. The GSK3B^KO^ cells were a generous gift from Dr. James R. Woodgett from the Lunenfeld-Tanenbaum Research Institute. We thank Jennifer Shellenberger for providing experimental and technical support in this study.

## Conflicts of Interest

The authors declare no conflict of interest.

